# Calbindin regulates Kv4.1 trafficking and excitability of dentate granule cells via CaMKII-dependent phosphorylation

**DOI:** 10.1101/2021.02.26.433035

**Authors:** Kyung-Ran Kim, Hyeon-Ju Jeong, Yoonsub Kim, Seung Yeon Lee, Yujin Kim, Hyun-Ji Kim, Suk-Ho Lee, Hana Cho, Jong-Sun Kang, Won-Kyung Ho

## Abstract

Calbindin, a major Ca^2+^ buffer in dentate granule cells (GCs), plays a critical role in shaping Ca^2+^ signals, yet how it regulates neuronal functions remains largely unknown. Here, we found that calbindin knock-out mice (CBKO) exhibited hyperexcitability in dentate GCs and impaired pattern separation, which was concurrent with reduced K^+^ current due to downregulated surface expression of Kv4.1. Consistently, manipulation of the calbindin expression in HT22 led to changes in CaMKII activation and the level of surface localization of Kv4.1 through phosphorylation at serine 555, confirming the mechanism underlying neuronal hyperexcitability in CBKO. We also discovered that Ca^2+^ buffering capacity was significantly reduced in the GCs of Tg2576 to the level of CBKO GCs, and this reduction was restored to normal levels by antioxidants, suggesting that calbindin is a target of oxidative stress. Our data suggest that regulation of CaMKII signaling by Ca^2+^ buffer is crucial for neuronal excitability regulation.

## Introduction

Ca^2+^ signaling is involved in every aspect of neuronal functions, from normal physiology, such as synaptic transmission and memory formation, to the pathogenesis underlying various neurologic and psychiatric diseases. Cytosolic Ca^2+^ buffers are essential components for the maintenance of Ca^2+^ homeostasis and the shaping of Ca^2+^ signals (Schwaller, 2012). Calbindin-D_28k_ (CB) is a major Ca^2+^ buffer of mature granule cells (GCs) in the dentate gyrus (DG) of the hippocampus (Celio, 1990), where half of the Ca^2+^ buffering is attributable to CB (Lee et al., 2009). CB is typically a mobile fast Ca^2+^ buffer, contributing to the shaping of the spatiotemporal extent of Ca^2+^ signals (Blatow et al., 2003). Increased CB levels by nerve growth factor (Iacopino et al., 1992b) and a lack of CB in degenerated substantial nigra (Iacopino et al., 1992a) suggest that CB plays a protective role from excitotoxicity. Interestingly, the expression of CB in the DG is markedly reduced in various pathologic conditions that are accompanied by cognitive dysfunction, including Alzheimer’s disease (AD) (Iacopino and Christakos, 1990; Palop et al., 2003; Stefanits et al., 2014) and schizophrenia/bipolar disorders (Altar et al., 2005; Walton et al., 2012; Yamasaki et al., 2008). Downregulation of CB in hippocampal excitatory neurons by early life stress is implicated in increasing susceptibility to stress-induced memory deficits (Li et al., 2017). However, a mechanistic understanding of the pathophysiologic roles of CB reduction is poor.

Ca^2+^/calmodulin (CaM)-dependent protein kinase II (CaMKII) is a multifunctional serine/threonine-protein kinase that is highly concentrated in the brain, and activity-dependent activation of CaMKII plays a central role in synaptic plasticity (Bayer and Schulman, 2019) and neuronal excitability (Hund et al., 2010; Zybura et al., 2020). Abnormal CaMKII activity has been implicated in various neuronal diseases, including schizophrenia (Yamasaki et al., 2008), intellectual disability (Küry et al., 2017), and AD (Reese et al., 2011), highlighting the importance of CaMKII in neuronal functions Calcineurin, a Ca^2+^-dependent protein phosphatase, plays a critical role in the activity-dependent alteration of synaptic transmission (Winder and Sweatt, 2001). It is intriguing to know Ca^2+^ buffers affect Ca^2+^-dependent protein kinase and phosphatase activities, thereby regulating neuronal functions. Overexpression of CB was found to promote neuronal differentiation, which was concurrent with CaMKII activation and inhibited by a CaMKII inhibitor (Kim et al., 2006), suggesting a link between CaMKII and CB in hippocampal progenitor cells. It is of great interest whether the interplay between CB and CaMKII plays a role in mature neurons.

The DG of the hippocampus has long been postulated to mediate pattern separation by transforming similar inputs into distinct neural representations (Leutgeb et al., 2007; O’Reilly and McClelland, 1994; Treves and Rolls, 1994). The sparse activity of GCs has been regarded as essential for the computational function to accomplish pattern separation (O’Reilly and McClelland, 1994; Rolls, 2013; Treves and Rolls, 1994). We have recently discovered that Kv4.1 is a key ion channel for the low-frequency firing of GCs, and mice with Kv4.1 depletion in the DG show impaired pattern separation (Kim et al., 2020), highlighting the role of Kv4.1 in DG functions. It is intriguing to investigate the regulatory mechanisms of Kv4.1.

In the present study, we discovered that the GCs in CB knock-out (CBKO) mice show increased neuronal excitability with reduced Kv4.1 activity and CaMKII activation. CB deficiency results in impaired CaMKII activation, which in turn reduces the surface localization of Kv4.1. CaMKII interacts with and phosphorylates Kv4.1 protein at serine 555 (S555), which regulates its targeting to the membrane and channel activities. Furthermore, CBKO mice showed impaired pattern separation. Our data collectively suggest that CB reduction predisposes patients to cognitive dysfunction, at least partially, by disrupting the CaMKII-dependent regulation of Kv4.1.

## RESULTS

### Increased excitability in the CBKO GCs

Reduced Ca^2+^ buffering may affect the Ca^2+^ homeostasis of cells. However, the relevance between the extent of Ca^2+^ buffering and the specific changes in neuronal functions is not well characterized. Using CBKO mice, we tested the possibility that Ca^2+^ dysregulation caused by reduced Ca^2+^ buffering could alter neuronal excitability in mature GCs of the DG, where CB is a major Ca^2+^ buffer (Lee et al., 2009). To avoid the effect of other Ca^2+^ buffers in neural progenitor cells and newborn immature neurons, we chose mature GCs located in the outer granular cell layer that has an input resistance (R_in_) lower than 200 MΩ. We confirmed that the average resting membrane potential (RMP; −81.6 ± 1.8 mV, *n* = 9, for control GCs; −84.4 ± 1.0 mV, *n* = 8 for CBKO GCs, p = 0.198) and R_in_ values (145.1 ± 11.4 MΩ for control GCs; 159.1 ± 13.4 MΩ for CBKO GCs, p = 0.439) were not significantly different between control and CBKO GCs. Interestingly, we found that the firing frequency in response to long depolarizing pulses was significantly higher in the CBKO GCs (red, Fig. 1Aa) than in the control GCs (black, Fig. 1Aa). As a measure of neuronal excitability, we plotted the firing frequencies (F) against the amplitude of the injected currents (I). The F-I relationship was steeper in CBKO (Fig. 1Ab), suggesting that abnormal Ca^2+^ homeostasis caused by reduced Ca^2+^ buffering induces ion channel remodeling in the GCs, which in turn induce hyperexcitability. The increased firing frequency was accompanied by a shortening of onset time of action potential (AP; 37.7 ± 1.8 ms, *n* = 8 for control; 23.9 ± 3.9 ms, *n* = 6 for CBKO GCs, p = 0.01388, Fig. 1Ba, Bb). AP threshold and AP shapes, measured by overshoot and half-width, were not affected in the CBKO GCs (Fig. 1Bb). These results suggest that the involvement of Na^+^ channels is less likely, but the downregulation of the K^+^ channel may cause the hyperexcitability of CBKO GCs. In contrast, no significant difference was found in CA1 pyramidal neurons between control and CBKO mice (Figs. 1Ca, Cb), suggesting that the changes in ion channels that cause hyperexcitability in CBKO are specific to GCs.

**Figure 1.**
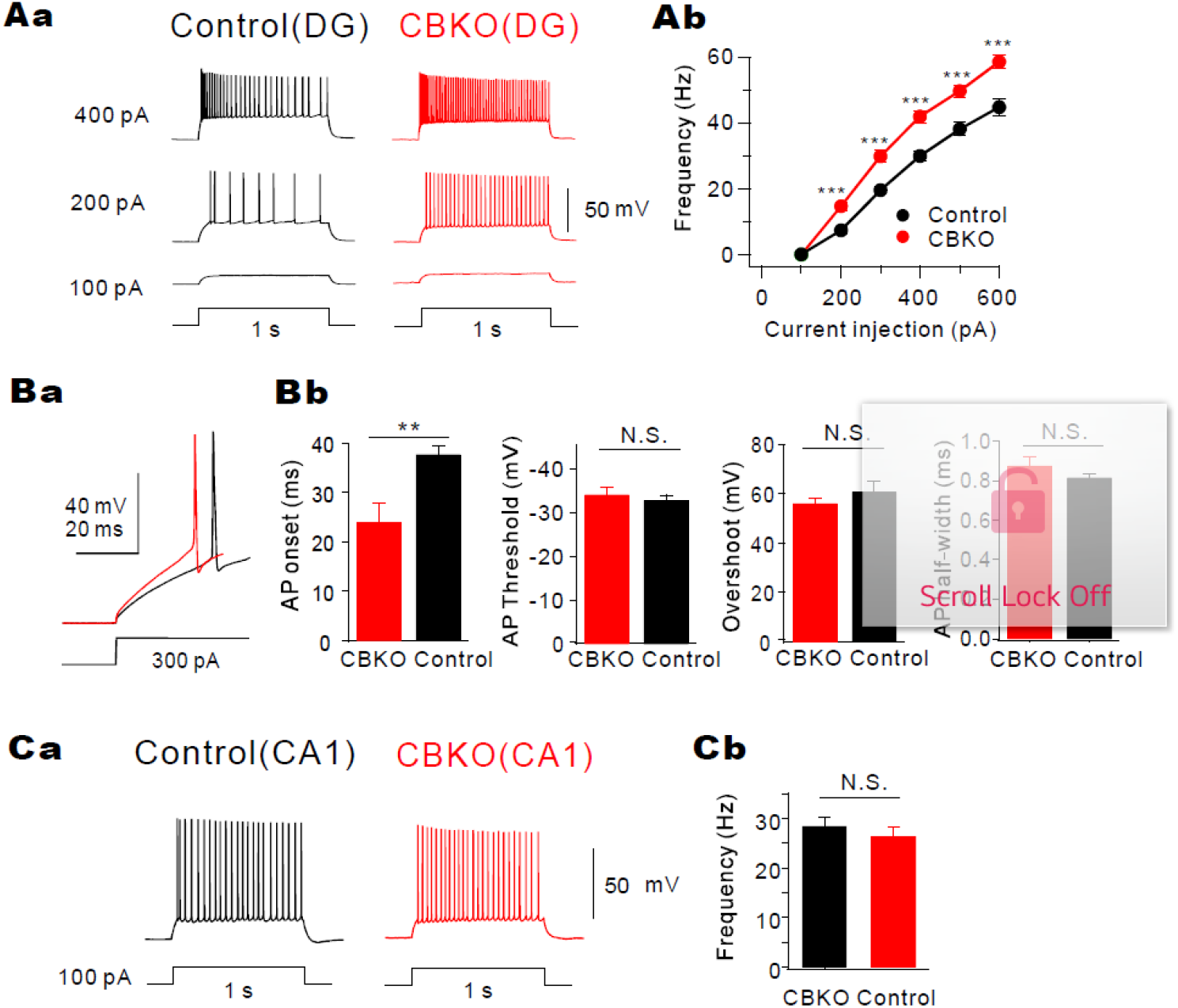
Increased firing frequency in dentate GCs of CBKO. ***Aa,*** Train of APs induced in dentate GCs of control (black) and CBKO mice (red) by 1 s depolarizing current injections as indicated by numbers left to the traces. ***Ab,*** The F-I curves in control (black, *n =* 9) and CBKO (red, *n =* 8). ***Ba***, Examplary traces of first AP elicited by 300 pA current injection from control (black) and CBKO (red) GCs. ***Bb***, The bar graphs showing the onset time, threshold, overshoot amplitude, and half-width of the first AP elicited by 300 pA current injection measured from CBKO (red) and control (black). ***Ca***, Trains of APs induced in CA1 of control (black) and CBKO (red) by 100 pA, 1 s depolarizing current injection. ***Cb,*** Bar graph for AP frequency at 100 pA current injection in CA1 cells of control and CBKO. Data are represented as mean ± SEM. **p < 0.01, ***p < 0.001, N.S. (not significant) p > 0.05 by Student’s t-test.

### Impaired pattern separation in CBKO mice

Low excitability of DG is considered important for information processing in the hippocampus, particularly for pattern separation (Rolls and Kesner, 2006). We addressed whether the increased firing of GCs in CBKO mice leads to an impairment in pattern separation during the contextual fear discrimination test. We first subjected CBKO mice to contextual fear conditioning using a pair of similar contexts (A and B). Both had the same metal grid floor, but B had a unique odor (1% acetic acid), dimmer lighting (50% of A), and a sloped floor at a 15° angle. As shown in Fig. 2A, the mice learned to discern a similar context (context B) over several days with a single footshock in context A. On the first three days, mice were placed only into A, receiving a single footshock after 180 s. On days four and five, the mice of each genotype were divided into two groups. One group of each genotype visited context A on day four and visited B on day five, and the other group visited B on day four and A on day five. No group received a footshock in either context, and freezing was evaluated for 5 min. Both genotypes showed identical freezing kinetics across the 5 min test in context A (Fig. 2Ba) and equivalent generalization between contexts (Fig. 2Bb, two-way ANOVA, genotype: *F*_(1,32)_ = 0.30, *p* = 0.58, context: *F*_(1,32)_ = 1.17, *p* = 0.29, genotype × context: *F*_(1,32)_ = 0.03, *p* = 0.86). The mice were subsequently trained to discriminate these contexts by visiting the two contexts daily for eight days in 2 h intervals (from day six to day 13), always receiving a footshock 180 s after being placed in context A but not in context B. The daily discrimination ratio was calculated as the ratio of freezing during the 180 s in context A to the total freezing during both visits (A and B). On day six, both genotypes could not distinguish the differences between contexts (Figs. 2Ca, Cb, genotype: *F*_(1,32)_ = 0.005, *p* = 0.94, context: *F*_(1,32)_ = 0.001, *p* = 0.97, genotype × context: *F*_(1,32)_ = 0.32, *p* = 0.57), so the discrimination ratio was about 0.5. As the experiment progressed, the control mice could discriminate context B from context A well, and the discrimination ratio increased. However, CBKO mice exhibited significant deficits in the acquisition of discrimination ability (Fig. 2Ca) and showed elevated freezing in the shock-free context B (Fig. 2Cb, t-test, Context B, *p* < 0.0001, two-way ANOVA, genotype: *F*_(1,32)_ = 6.71, *p* = 0.01, context: *F*_(1,32)_ = 52.99, *p* < 0.0001, genotype × context: *F*_(1,32)_ = 25.60, *p* < 0.0001). To examine whether impaired discrimination between similar contexts in CBKO mice is due to a problem with memory acquisition, we examined the context specificity of the conditioning by assessing freezing behavior using a distinct pair of contexts (A and C). This distinct context (context C) evoked significantly lower levels of freezing (similar in the two genotypes) than in context A (Figs. 2Da, Db, genotype: *F*_(1,11)_ = 0.002, *p* = 0.96, context: *F*_(1,11)_ = 30.34, p < 0.0001, genotype × context: *F*_(1,11)_ = 0.002, *p* = 0.96). These data imply that CBKO mice have no problem in learning or discriminating between very distinct objects but have difficulty in discriminating between similar contexts (pattern separation). We also tested the locomotor activity and anxiety levels of CBKO mice using the open field test. Control and CBKO mice showed similar exploratory patterns as they moved the same distance in the open field box and spent a similar amount of time in the center zone (Figs. 2Ea, Eb).

**Figure 2.**
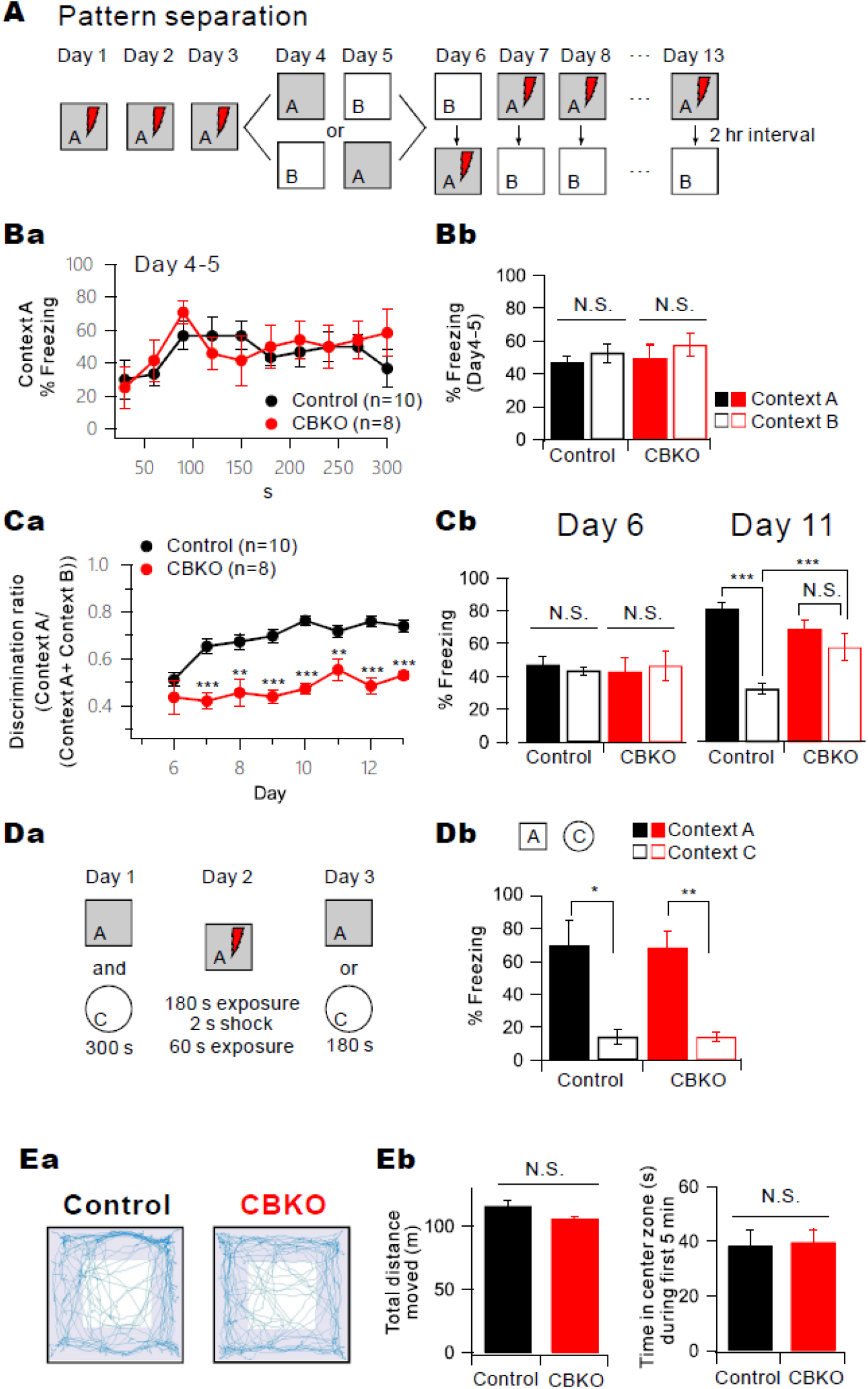
Impaired pattern separation in CBKO mice. *A*, Experimental procedure for pattern separation in 15- to 19-week-old control (*n* = 10) and CBKO (*n* = 8) mice. *Ba*, On Day 4 and 5, the kinetics of freezing across the 5 min test in context A. *Bb*, The percentage of freezing in context A (filled bar) and context B (open bar) during Day 4 to 5 in both contexts (A and B). Control (black, *n* = 10) and CBKO (red, *n* = 8) mice displayed equal amounts of freezing in both contexts (A and B). *Ca*, On Day 6 to 13, time course of the discrimination ratio in control (black, *n* = 10) and CBKO (red, *n* = 8) mice. *Cb*, The percentage of freezing in context A (filled bar) and context B (open bar) for the control (black, *n* = 10) and CBKO (red, *n* = 8) mice on Day 6 (left) and Day 11 (right). *Da*, Experimental procedure for one-trial contextual fear conditioning between control (*n* = 8) and CBKO (*n* = 8) mice. ***Db***, The percentage of freezing in context A (filled bar) and context C (open bar, distinct object) for the control (*n* = 8) and CBKO (*n* = 8) mice. ***Ea***, The trajectories traveled by the control (*left*) and CBKO (*right*) mice in the open field test showed how the mice traveled. ***Eb***, Summary bar graph of the total traveled distance (*left*) and time in center zone during first 5 min (*right*). Total distance moved (m), Control, n = 6, 116.20 ± 4.45, CBKO, n = 6, 106.11 ± 1.79, p = 0.06175; Time in center zone (sec), Control, n = 6, 11.81 ± 1.22, CBKO, n = 6, 13.91 ± 1.55, p = 0.31208. Data are represented as mean ± SEM.

### Reduced Kv4.1 current underlies increased excitability in the CBKO GCs

To understand the ion channel mechanisms responsible for the hyperexcitability in CBKO GCs, we analyzed the difference in K^+^ currents between CBKO and control CGs. Whole-cell K^+^ currents were recorded in the voltage-clamp mode by applying a depolarizing voltage step from −60 mV to +50 mV (in 10 mV increments, 1 s duration) from a holding potential of −70 mV. TTX, Cd^2+^/Ni^2+^, bicuculline, and CNQX were added to the external solution to block the Na^+^ channels, Ca^2+^ channels, GABA_A_ receptors, and AMPA/kainate receptors, respectively. Total K^+^ currents were dissected into TEA-sensitive (I_TEA_) and 4-AP-sensitive K^+^ currents (I_4-AP_) (Fig. 3A). I_TEA_ obtained from CBKO GCs (*n* = 9) showed no significant difference compared to I_TEA_ in control GCs reported previously (Kim et al., 2020) both with peak (I_peak_ at +30 mV; 1.807 ± 0.152 nA, *n* = 9, for control; 1.727 ± 0.096 nA, *n* = 9, for CBKO, p = 0.66) and steady-state currents (I_ss_ at +30 mV; 1.107 ± 0.126 nA, *n* = 9, for control: 1.012 ± 0.133 nA, *n* = 9, for CBKO, p = 0.609) (Figs. 3Bb, Bc). In contrast, the characteristics of I_4-AP_ in CBKO GCs were different from those in control GCs. The inactivation phase of I_4-AP_ in CBKO GCs showed a fast time constant with little I_ss_ (Fig. 3Ca), which are typical characteristics of A-type K^+^ current, while that in the control GCs had a significant proportion of I_ss_, which is attributable to Kv4.1 current (Fig. 3Cb) (Kim et al., 2020). The amplitude of I_4-AP_ in CBKO GCs was significantly smaller than that in the control GCs (Figs. 3Cc). The reduction in I_ss_ (63%) was more pronounced than the reduction of I_peak_ (36%). I_4-AP_ in CBKO GCs was similar to I_4-AP_ in the control GCs in the presence of Kv4.1 antibody, as reported previously (Kim et al., 2020). This suggests that the outward K^+^ current component reduced in the CBKO GCs is likely to be the Kv4.1 current.

**Figure 3.**
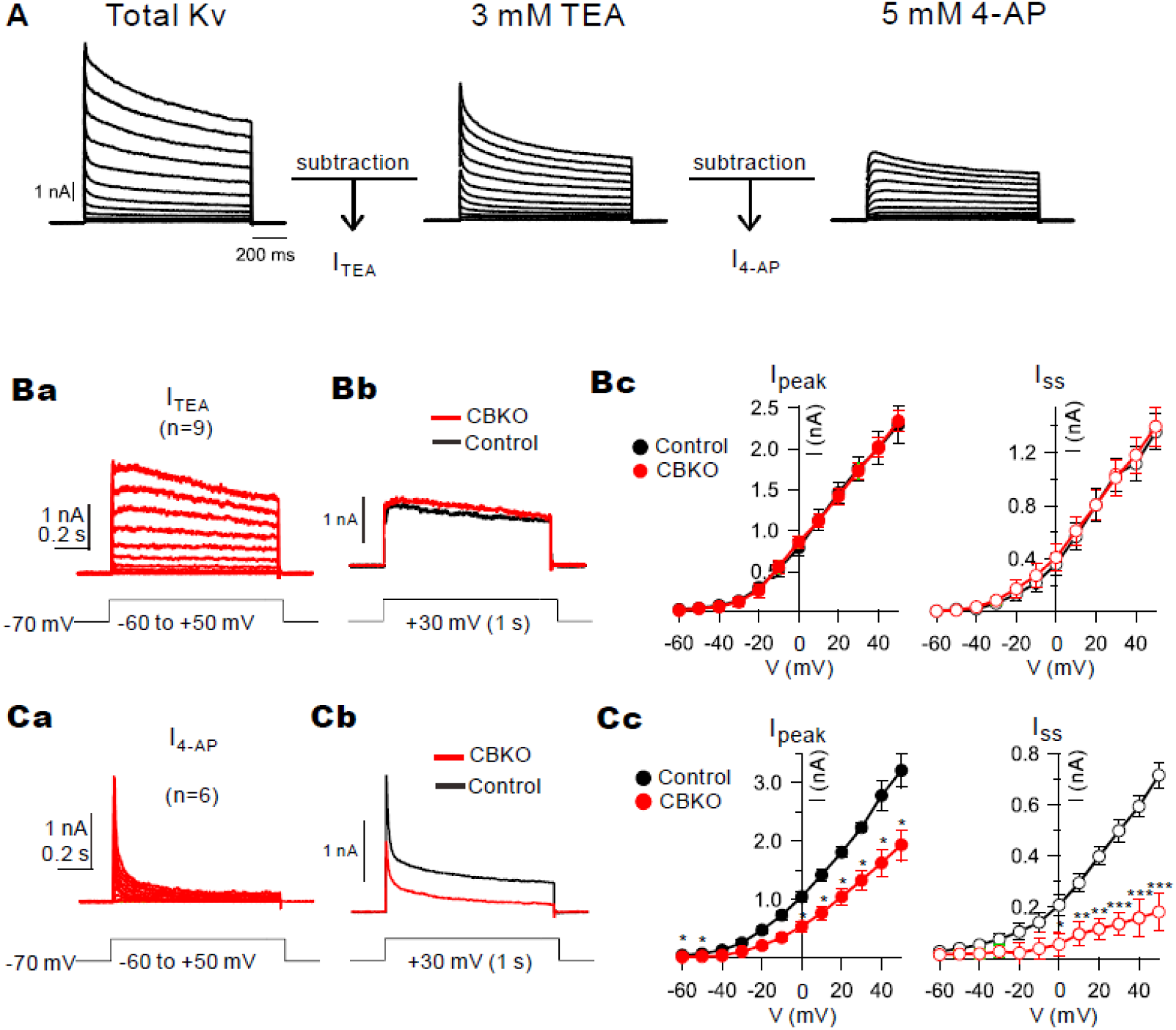
Decrease 4-AP-sensitive current in CBKO GCs. *A*, Representative whole-cell, voltage-gated K^+^ currents from control dentate GCs. Currents were evoked in response to 1 s, 10 mV voltage steps to potentials between −60 mV to +50 mV from a holding potential of −70 mV. After recording the total outward currents, we changed the bath solution which contained 3 mM TEA, and the difference currents before and after TEA were measured to obtain TEA-sensitive currents (I_TEA_). Then 5 mM 4-AP was added to obtain 4-AP sensitive currents (I_4-AP_) sequentially. *Ba*, I_TEA_ obtained from CBKO GCs (*n = 9*) were averaged. *Bb,* Superimposed I_TEA_ traces at +30 mV measured CBKO (red) and control (black) at +30 mV. *Bc*, Amplitudes of the peak currents (I_peak_) and steady state currents (I_ss_) for I_TEA_ are plotted as a function of the given potential (V). Control (black closed circle, *n = 4*) and CBKO (red closed square, *n = 9*). *Ca,* The difference currents before and after applying 5 mM 4-AP. *Cb,* Superimposed I_4-AP_ traces measured from CBKO (red) and control (black) at +30 mV. *Cc*, I_peak_ and I_ss_ for 5 mM 4-AP sensitive I_A_ currents same as (*Bc)*. Control (black closed circle*, n = 9*) and CBKO (red closed circle, *n = 6*). Data are represented as mean ± SEM. *p < 0.05, **p < 0.01, ***p < 0.001, N.S. not significant (p > 0.05) by Student’s t-test.

### Calbindin regulates cell surface localization of Kv4.1 via CaMKII-mediated phosphorylation

To further confirm that the K^+^ current component reduced in CBKO GCs is the Kv4.1 current, we monitored changes in the outward current amplitude during perfusion of the Kv4.1 antibody through a patch pipette after patch break-in. Kv4.1 antibody significantly reduced the outward K^+^ currents in the control mice but did not induce changes in CBKO GCs (Figs. 4Aa, Ab), suggesting that Kv4.1 is not functional in CBKO GCs.

**Figure 4.**
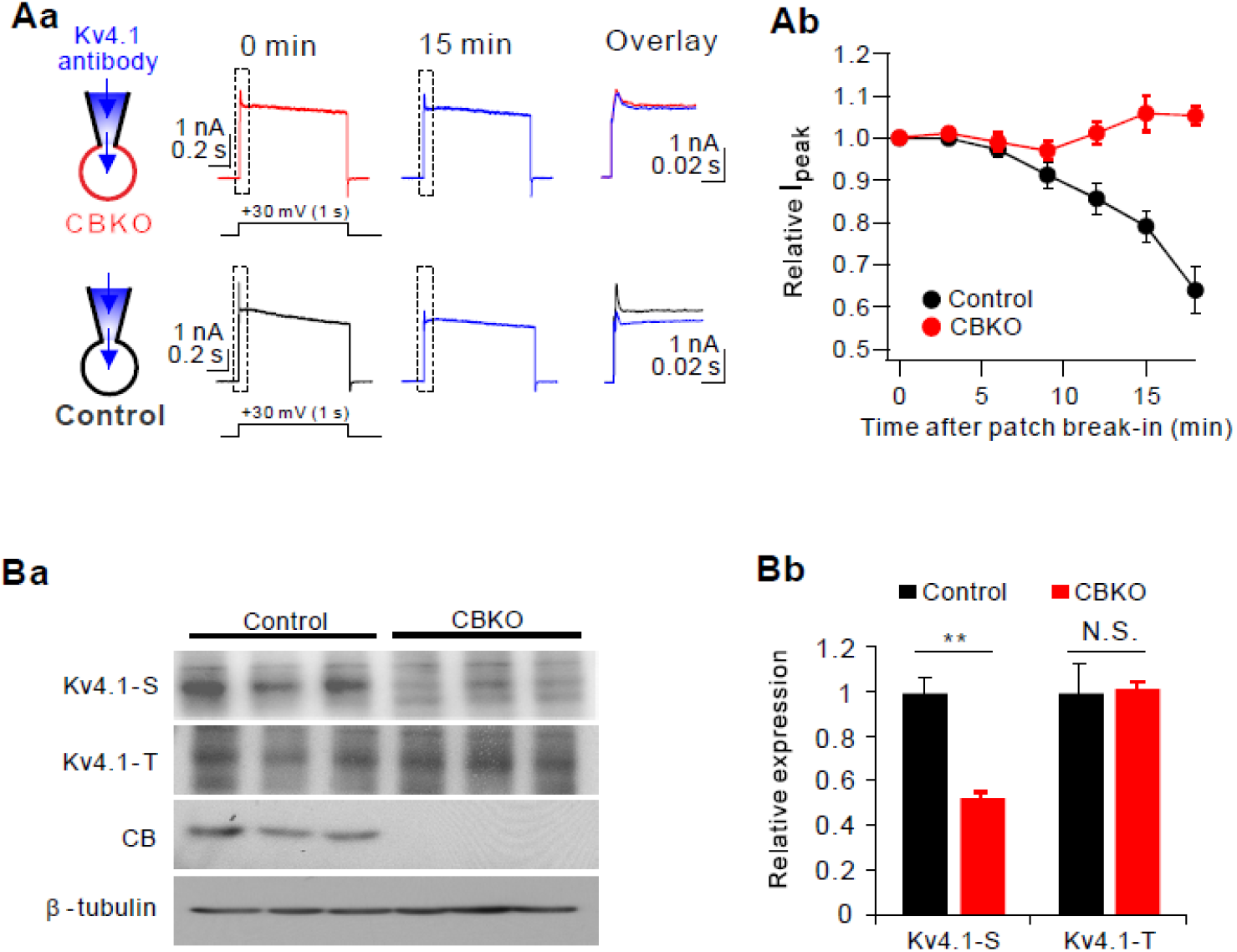
Evidence for the reduced Kv4.1 currents in CBKO GCs. ***Aa,*** Top: Whole-cell, voltage-gated K currents activated by + 30 mV depolarizing pulses in CBKO GCs with intracellular solution containing Kv4.1 antibody recorded shortly after ruptured (0 min) and fully diffused (15 min). Each trace is expanded and superimposed for comparison (dashed box). Bottom: Same as described at (top) but in control GCs. ***Ab,*** Time course of normalized I_peak_ at each time point after break-in. ***Ba,*** Western blot analysis for total Kv4.1 (Kv4.1-T) and surface Kv4.1 (Kv4.1-S) in DG of 8-week-old control and CBKO mice. Absence of calbindin (CB) expression was confirmed in DG of CBKO mice. For a loading control, β-tubulin was used. ***Bb,*** Relative protein levels of Kv4.1 in surface or total lysates in CBKO compared to control. Values are means of three determinants ± SD. (*n* = 3). **p < 0.01, N.S. not significant by Student’s t-test.

To investigate the mechanism of reduced Kv4.1 current in CBKO GCs, we first tested alterations in Kv4.1 protein expression and surface localization. To monitor membrane localization, isolated DG was labeled with biotin, lysed, and subjected to pulldown with streptavidin-beads. The immunoblot analysis revealed that the level of the surface-resident Kv4.1 (Kv4.1-S) was significantly reduced in the CBKO DG, while the level of total Kv4.1 protein (Kv4.1-T) was not significantly altered relative to the control DG (Figs. 4Ba, Bb). These results suggest that the reduced Kv4.1 current in CBKO GCs are attributable to the impairment of Kv4.1 trafficking to the surface membrane.

To further understand the mechanism underlying changes in Kv4.1 trafficking by reduced CB, we infected a mouse hippocampal neuronal cell line HT22 with lentiviruses expressing either control scrambled or CB-shRNA and examined the surface expression of Kv4.1 in differentiated cells. Similar to the results obtained with the CBKO DG, the level of membrane-resident Kv4.1 protein was specifically reduced in CB-depleted HT22 cells without any reduction in the total Kv4.1 level (Fig. 5A). We then examined the level of active phosphorylated CaMKII α (p-CaMKIIα), which acts as the main player in the Ca^2+^ signaling pathway implicated in diverse cellular processes, including the control of neuronal excitability (Ohno et al., 2006). The level of p-CaMKII α was substantially reduced in CB-depleted HT22 cells (Fig. 5A). Conversely, CB overexpression enhanced the levels of surface Kv4.1 and p-CaMKllα, relative to the control (Fig. 5B). These data suggest that CaMKIIα might be a downstream effector of CB to control Kv4.1 surface localization, likely through phosphorylation. Thus, we investigated the potential phosphorylation of Kv4.1 in relation to the CB levels and CaMKII activities. HEK293T cells were transfected with the expression vectors for Kv4.1-GFP with or without CB, and two days later, the cells were treated with a CaMKII inhibitor KN93 or the relevant control compound KN92 for 24 h. To monitor the phosphorylated Kv4.1 level, cell lysates were subjected to immunoprecipitation with anti-phospho-serine antibodies and immunoblotting analysis. Co-expression of CB and Kv4.1 significantly elevated the phosphorylated Kv4.1 protein level in the treatment with control KN92, which was abrogated by KN93 treatment (Fig. 5C). These data suggest that CB facilitates the CaMKII-mediated phosphorylation of Kv4.1, thereby increasing its cell surface expression in the DG and hippocampal cells.

**Figure 5.**
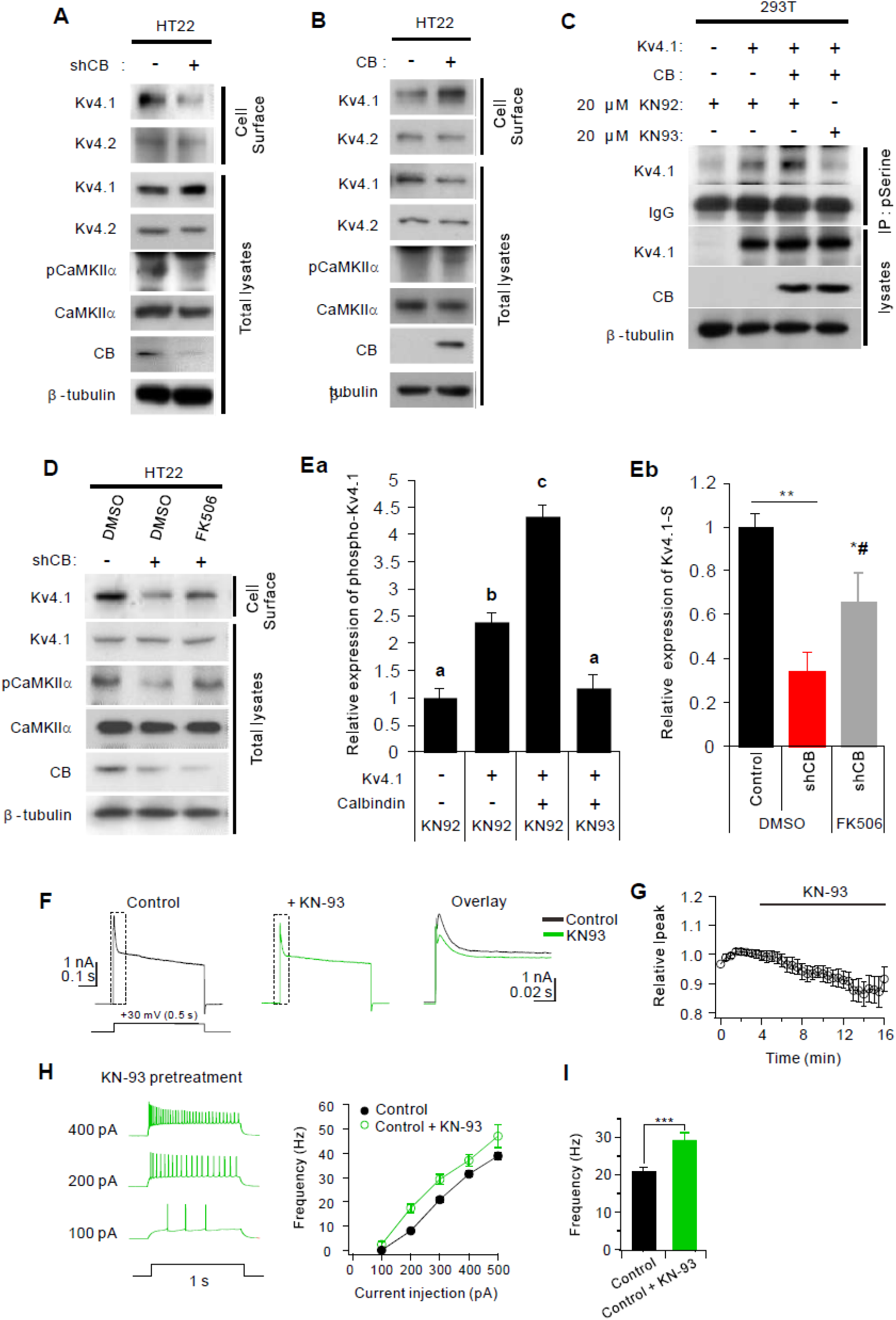
Calbindin regulates cell surface localization of Kv4.1 via CaMKII-mediated phosphorylation. ***A, B,*** Cell-surface biotinylation assay of control and shRNA Calbindin (A) or Calbindin (B) transfected HT22 cells followed by the immunoblot analysis with indicated antibodies. ***C***, Analysis of serine-phosphorylation of Kv4.1 in 293T cells expressing control or Kv4.1 and Calbindin treated with 20 μM KN92 or KN93 for 24 hours. ***D***, Surface biotinylation assay of control and Calbindin-depleted HT22cells treated with vehicle or 10 μM FK506 for 1 hour followed by the immunoblot analysis with indicated antibodies. ***Ea***, Quantification for relative phosphorylation levels of Kv4.1 at serine residues shown in panel C. The intensities were normalized to the level of β-tubulin. Values are means of triplicate determinants (ANOVA Tukey). Letters indicate statistically distinct groups (p < 0.05). ***Eb***, Quantification for relative cell-surface protein levels of Kv4.1 shown in panel D. The intensities were normalized to the level of β-tubulin. *p < 0.05, **p < 0.01 versus control group; #p < 0.05 versus shCB group. ***F,*** Whole-cell K^+^ currents at + 30 mV in control GCs before (black) and after KN-93 application (green). ***G,*** Time course of the changes in K^+^ current amplitude after KN93 application (average from 7 cells). ***H,*** Representative AP traces induced by injecting depolarizing currents as indicated by number left to the traces in control GCs pretreated in KN-93 (*left*), and corresponding F-I curve (green circles, *right*). F-I curve from control GCs shown in Fig. 1Ab was superimposed for comparison (black line) ***I,*** Firing frequency obtained by 200 pA current injection obtained in the presence of KN-93 in control GCs (green, *n = 7*) is compared with that in control (black) shown in Fig. 1A. Data are represented as mean ± SEM. ***p < 0.001. N.S. not significant by Student’s t-test.

Since the Ca^2+^ buffering capacity of CBKO GCs is only half that of control GCs (Lee et al., 2009), the intracellular Ca^2+^ concentration is expected to be higher in CBKO GCs (CBKO GCs are subject to Ca^2+^ overload). Thus, it appears to be paradoxical that CaMKII inhibition shows a phenotype similar to that of CBKO. To understand how p-CaMKII is downregulated by CB knockdown, we analyzed the effect of a Ca^2+^-dependent phosphatase inhibitor, FK506, on p-CaMKII and the surface localization of Kv4.1 in CB-depleted HT22 cells (Figs. 5D and E). CB depletion reduced the level of surface Kv4.1 protein in the treatment with vehicle dimethyl sulfoxide (DMSO), while FK506 treatment significantly restored the surface Kv4.1 levels (Figs. 5D and E). These results suggest that CB knockdown may activate Ca^2+^-dependent phosphatase, which in turn decreases p-CaMKII levels and Kv4.1 surface localization.

To confirm whether CaMKII-dependent phosphorylation is a key mechanism for Kv4.1 activity in GCs, we examined the effect of CaMKII inhibitors (KN93 and KN62) on outward K^+^ currents (Figs. 5F, G, S1). In control GCs, outward K^+^ currents were reduced significantly by KN93 (625.6 ± 160.9 pA at +30 mV, *n* = 7), which was comparable to the Kv4.1 current amplitude in control GCs reported previously (619.6 ± 66.4 pA) (Kim et al., 2020). We then obtained I_4-AP_ from GCs that were pretreated with KN62 for more than 20 min (Fig. S1A). As we previously reported (Kim et al., 2020) and shown in Fig. 3Cb, a characteristic feature of I_4-AP_ in control GCs is a relatively large sustained current (I_ss_), which is attributable to Kv4.1 current, but I_4-AP_ in the presence of KN62 had a small I_ss_. For comparison, we superimposed I_4-AP_ at +30 mV obtained in the presence of KN62 with that obtained from control GCs and CBKO GCs. This shows that I_4-AP_ in the presence of KN62 was significantly smaller than that in control GCs but comparable to that in CBKO (Fig. S1B). Accordingly, GCs pretreated with CaMKII inhibitors displayed hyperexcitability, which was comparable to that of CBKO GCs (Figs. 5H, I). These results support that the outward K^+^ current component decreased by CaMKII inhibitors is also reduced in CBKO GCs. In contrast, KN93 did not show a significant effect in CBKO GCs (Fig. S2), which was consistent with the idea that the Kv4.1 current that requires CaMKII is not functional in CBKO GCs.

### CaMKII regulates Kv 4.1 activities by phosphorylating S555 of Kv4.1

A sequence analysis to predict the phosphorylation sites in Kv4.1 by CaMKII using the PhosphoSite database revealed three potential serine residues at 265, 555, and 568 of Kv4.1 (Fig. 6A). Thus, we generated serine to alanine mutants for these sites in Kv4.1 and analyzed the phosphorylation status by CaMKII activities. Co-transfection of Kv4.1 with CaMKIIα elevated Kv4.1 levels immunoprecipitated with anti-phosphoserine antibody, compared to the control Kv4.1 single expression (Fig. 6B, C). The phosphorylated levels of R265A and S568A mutants were equivalent to wild-type Kv4.1 in response to CaMKIIα. In contrast, the S555A mutant failed to show any increase in response to CaMKIIα, suggesting that S555 is a target phosphorylation site for CaMKII α.

**Figure 6.**
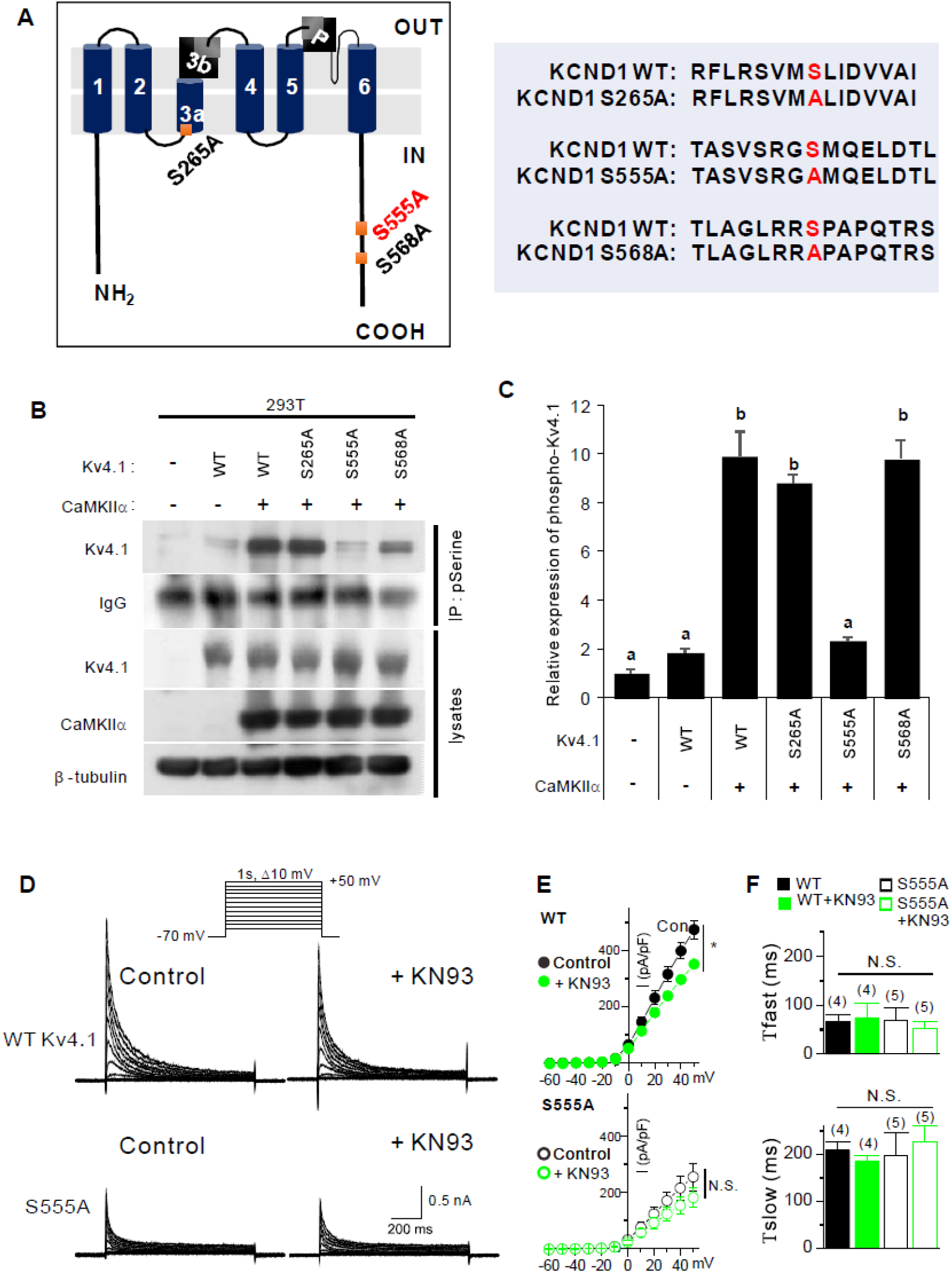
CaMKII regulates Kv 4.1 activities by phosphorylating S555 of Kv4.1. ***A***, Schematic representation of the domain structure of Kv4.1. Potential serine-phosphorylation sites are labeled red. (left) Alignments of the potential serine-phosphorylation site are shown in red and serine residue (S) that are switched to alanine (A). ***B***, Immunoprecipitation with anti-phosphoserine antibody and immunoblotting with indicated antibodies in 293T cells expressing control, WT Kv4.1, or Mutant Kv4.1 (S265A, S555A, and S568A) with CaMKll. ***C***, Quantification of serine phosphorylation levels of Kv4.1. Values are means of triplicate determinants (ANOVA Tukey, p < 0.05). Letters indicate statistically distinct groups. *D*, Representative current recordings from HEK293T cells expressing WT Kv4.1 (upper) or S555A Kv4.1 (lower) before and after KN93 (1 μM) treatment for 5 min. Currents were elicited by voltage steps from −60 mV to +50 mV at a holding potential of −70 mV. *E*, Current–voltage curves for peak current amplitude of WT and S555A Kv4.1 currents in the absence and presence of KN93 (1 μM). Data represent mean ± S.E.M. *P < 0.05, Paired Student’s t-test. *F*, The effect of S555A mutation and KN93 on the fast and slow inactivation time course. The inactivation time course of Kv4.1 currents were well fitted with a double-exponential function. Data are represented as mean ± SEM. N.S. not significant by Student’s t-test.

We further examined the correlation between CaMKII-induced phosphorylation and Kv4.1 channel function by recording whole-cell currents in HEK293T cells expressing WT or S555A Kv4.1 (Figs. 6D and E). Compared to WT, S555A Kv4.1 exhibited remarkably decreased currents. The current density at +50 mV was 473.58 ± 32.3 pA/pF (*n* = 4) for WT Kv4.1, and it decreased to 254.86 ± 49.34 pA/pF (*n* = 5; P<0.05) for S555A Kv4.1. Furthermore, CaMKlla inhibition with KN93 decreased WT Kv4.1 activity to 350.36 ± 10.57 pA/pF (*n* = 4, p < 0.05 vs. WT Kv4.1 under control condition), while it could not further reduce the S555A Kv4.1 activity (180.92 ± 34.71 pA/pF, *n* = 5; p > 0.05 vs. S555A Kv4.1 under control conditions). The S555A mutation and KN93 treatment did not change the inactivation kinetics (Fig. 6F). These data suggest that the phosphorylation of the S555 residue in Kv4.1 by CaMKII is a key regulatory step in Kv4.1 activity.

### Functional deficit of calcium buffering in Tg2576 GCs is restored by antioxidant

Neuronal depletion of CB was reported to be tightly linked to AD-related cognitive deficits. However, this depletion appeared in at least six-month-old AD model mice (Palop et al., 2003). We recently reported that Kv4.1 is downregulated, leading to hyperexcitability in dentate GCs in one to two-month-old Tg2576 (Kim et al., 2021). Therefore, we asked whether dysfunctional Ca^2+^ buffering has pathophysiological significance in early preclinical stages. To this end, we measured the cellular Ca^2+^ buffer capacity in mature GCs from one to two-month-old WT and Tg2576 mice using a method described in our previous paper (Lee et al., 2009). To estimate κ_E_ (sum of the Ca^2+^ binding ratio of mobile and static endogenous Ca^2+^ buffers), we loaded GCs with fura-2 (250 μM) with whole-cell patch pipettes (R_S_ = 20.5 ± 1.0 MΩ), and evoked Ca^2+^ transients (CaTs) by applying voltage pulses (from −70 mV to 0 mV, 40 ms duration) at different time points of the fura-2 loading (Fig. 7A). As the cytosolic fura-2 concentration increased gradually after patch break-in, CaTs exhibited decreased amplitudes (*A*) and increased decay time constants (τ). To estimate the κ_E_ value, we plotted τ against κ_B_ and obtained the *x*-intercept from the extrapolation of the linear fit (Fig. 7B). The mean κ_E_ value in WT GCs estimated from the τ *vs* κ_B_ plots was 340.7 ± 12.5 (*n* = 10), while that in Tg2576 GCs was 206.8 ± 4.7 (n = 7), which was significantly lower than that in WT GCs (P = 0.00000034, Fig. 8C). The mean κ_E_ values estimated from the *A*^−1^ *vs* κ_B_ plots showed similar results (WT GCs: 342.4 ± 15.4, *n* = 10, Tg2576 GCs: 211.4 ± 19.4, *n* = 7, P = 0.000081, Fig. 7C). Notably, the κ_E_ value estimated in Tg2576 GCs is comparable to that in CBKO GCs reported previously (210.9 ± 4.8, red dashed line, Fig. 7C) (Lee et al., 2009). These results suggest that reduced Ca^2+^ buffering in Tg2576 GCs may be attributable to impaired CB function.

**Figure 7.**
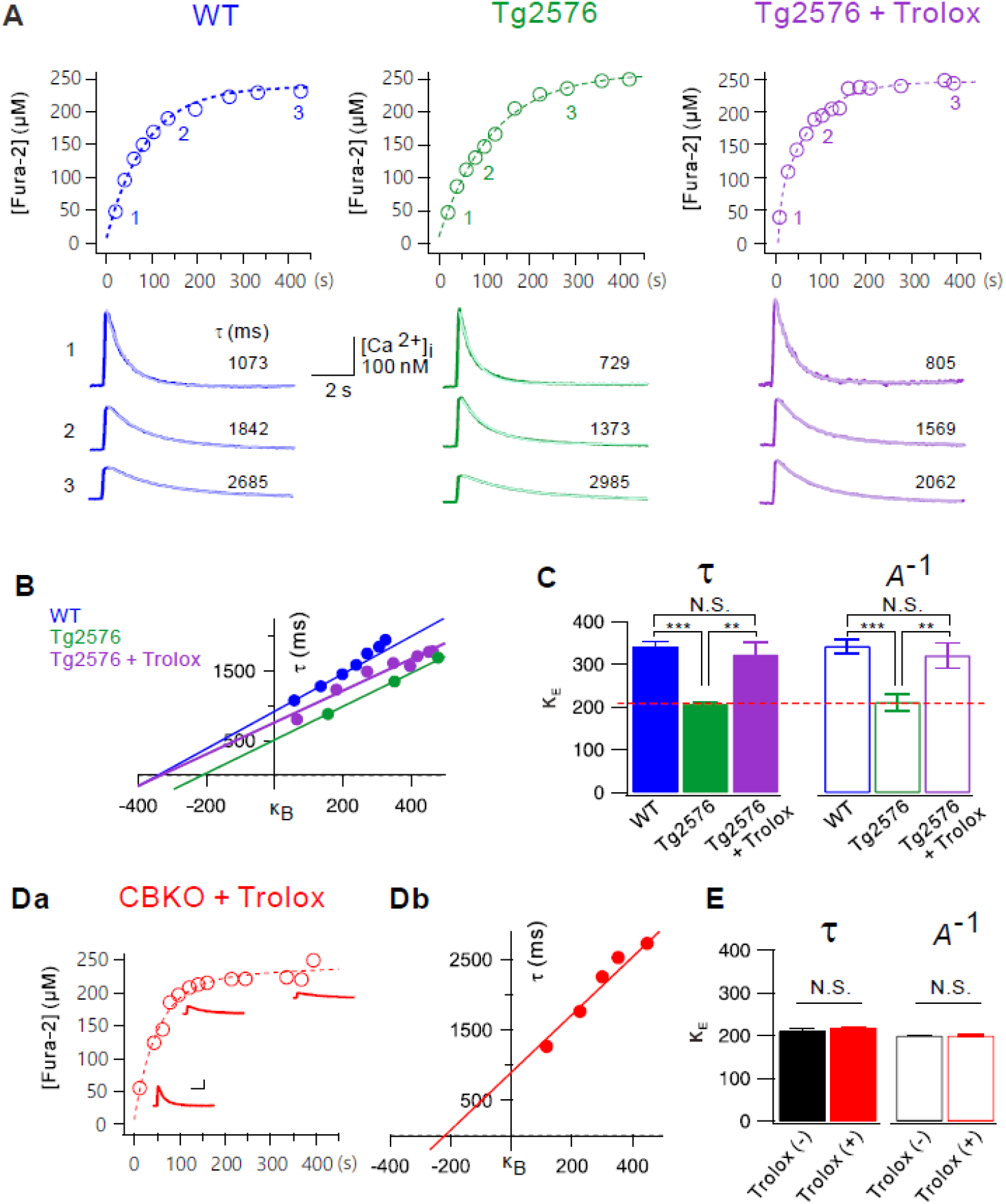
Endogenous Ca^2+^ binding ratios (κE) is reduced in dentate GCs of Tg2576 GCs. ***A***, Time course of Fura-2 loading (250 μM) during whole-cell patch recording in wild-type (WT, *blue*), Tg2576 (*green*), and Tg2576 slices pretreated with Trolox (500 μM, *violet*) for 1 hr. Ca^2+^ transients (CaTs) were evoked by depolarizing pulse (from −70 mV to 0 mV, 40 ms in duration). Fura-2 concentrations were calculated from isosbestic fluorescence and plotted as a function of whole-cell recording time. Exemplary CaTs shown underneath the loading curve were obtained at 20 s, 135 s and 427 s in WT, 40 s, 124 s and 282 s in Tg2576, and 9 s, 103 s and 392 s in Tg2576 + Trolox. Inset scale bars indicate 100 nM [Ca^2+^]_i_ and 2 s. ***B***, Plots of time constants (τ) for the decay phases of CaTs as a function of incremental Ca^2+^ binding ratios of fura-2 (κ_B_). An *x*-axis intercept of the linear fit to this plot was considered as the endogenous Ca^2+^ binding ratios (κ_E_). ***C***, Mean values for κ_E_ from WT (*blue, n=10*), Tg2576 (*green, n=7*) and Tg2576 + Trolox (*violet*, *n = 6*) dentate GCs estimated from the plots of τ (solid bars). The κ_E_ values were also estimated from the plot of 1/Amplitude versus κ_B_ (open bars). Dashed line (red) indicates the κ_E_ value of CBKO GCs reported previously (Lee et al., 2009). Data are represented as mean ± SEM. **p < 0.01, ***p < 0.001, N.S. not significant by Student’s t-test.

CB has two N-terminal cysteine residues that undergo redox-driven structural changes, and oxidized CB has a lower Ca^2+^ binding affinity than that of reduced CB (Cedervall et al., 2005). We previously showed that mitochondrial reactive oxygen species (ROS) production is increased in the dentate GCs of one to two-month-old Tg2576 mice, leading to depolarization of the mitochondrial potential, which was reversed by the antioxidant Trolox (Lee et al., 2012). To test the possibility that oxidative stress induces the oxidation of CB to compromise the Ca^2+^ buffering capacity, we examined whether the reduced κ_E_ in Tg2576 GCs was restored by antioxidant treatment. When hippocampal slices obtained from Tg2576 mice were incubated in Trolox (500 μM) for 1 h, the κ_E_ value of Tg2576 GCs was 322.98 ± 30.2 (*violet*, estimated from the τ *vs* κ_B_ plots, Fig. 7A), which was not significantly different from the κ_E_ value of WT GCs (*blue*, Fig. 7A, p = 0.53941). Reduced Ca^2+^ buffering and its restoration by Trolox in Tg2576 GCs suggest that CB function is impaired by oxidative stress.

## Discussion

The roles of endogenous Ca^2+^ buffers in the brain have been investigated mostly by analyzing the phenotype of mice either lacking or over-expressing one of the Ca^2+^ buffers (Schwaller, 2012). Mice with reduced CB expression show deficits in memory and hippocampal long-term potentiation (Molinari et al., 1996), reduced short-term facilitation (Yang et al., 2020), and impaired motor coordination (Airaksinen et al., 1997; Schwaller, 2012). In contrast, the overexpression of CB in DG GCs was shown to alter mossy fiber presynaptic function and impair hippocampal-dependent memory (Dumas et al., 2004). These data suggest that proper levels of CB are critical for neuronal activity control. However, the mechanisms underlying the involvement of CB in these phenotypes are largely unknown. In the present study, we investigated the cellular and molecular mechanisms underlying the excitability changes in dentate GCs in CBKO mice, and found that Ca^2+^ dysregulation caused by the loss of CB induces Kv4.1 downregulation, leading to hyperexcitability in dentate GCs. In addition, we showed impaired pattern separation in CBKO mice. These results, together with previous results that showed that mice selectively depleted of Kv4.1 in DG exhibit impaired pattern separation (Kim et al., 2020), highlighting the importance of the low excitability of dentate GCs in pattern separation.

CB loss in dentate GCs has been shown to be tightly linked to AD-related cognitive deficits (Palop et al., 2003). CB loss is also a well-known feature of temporal lobe epilepsy (Carter et al., 2008; Maglóczky et al., 1997; Nägerl et al., 2000). Since neuronal hyperactivity is also associated with AD, it has been suggested that CB loss is a consequence of neuronal hyperactivity. However, the possibility that CB loss is not merely an epiphenomenon of these diseases but causally related to DG dysfunction remains to be elucidated. In this respect, CBKO mice can be a useful model for studying the pathophysiological significance of CB loss in various diseases. In the present study, we proposed a causal link between Ca^2+^ buffering deficit and hyperexcitability of DG, which leads to cognitive deficits. Furthermore, we identified the molecular mechanism by which Ca^2+^ buffering deficits lead to neuronal hyperexcitability. We showed that CB depletion or overexpression in HT22 cells decreased or increased the level of surface Kv4.1 along with p-CaMKII levels (Figs. 5A, B). Furthermore, pharmacological inhibition of CaMKII reduced Kv4.1 surface expression (Fig. 5C), suggesting that Kv4.1 trafficking to the surface membrane is regulated by CaMKII-dependent phosphorylation. We further identified a phosphorylation site in Kv4.1, which is critical for channel trafficking (Fig. 6). However, it is unclear how reduced Ca^2+^ buffering inhibited CaMKII activity since reduced Ca^2+^ buffering was expected to increase cytosolic Ca^2+^ levels, and CaMKII activity was expected to increase in high Ca^2+^. To investigate this, we focused on another Ca^2+^-dependent enzyme that can act in the opposite way to kinase, calcineurin, a Ca^2+^-dependent phosphatase, and confirmed that reduced CaMKII activity and reduced Kv4.1 surface expression were restored, at least in part, with the pharmacological inhibition of calcineurin (Fig. 5D). These results suggest that Ca^2+^ dysregulation caused by CBKO resulted in an abnormal balance between Ca^2+^-dependent protein phosphatase and Ca^2+^-dependent protein kinase, causing reduced Kv4.1 phosphorylation, which is crucial for the surface localization of Kv4.1.

Protein phosphorylation and dephosphorylation are two essential mechanisms that regulate many functional proteins, such as enzymes, receptors, and ion channels. In the brain, Ca^2+^-dependent kinases and phosphatases play key roles in the activity-dependent regulation of neuronal functions. In particular, their roles in synaptic plasticity are well-known. A small increase in Ca^2+^ activates phosphatase, leading to long-term depression, while a higher increase in Ca^2+^ activates kinases, leading to long-term potentiation (Lisman, 1989). These results imply that Ca^2+^ signals are translated into the balance between Ca^2+^-dependent kinases and phosphatases, thus determining the phosphorylation states of synaptic proteins to regulate synaptic plasticity. Activity-dependent regulation of ion channels through the balancing of Ca^2+^-dependent kinases and phosphatases has also been well documented. Activity-dependent downregulation of Kv1.2 activity in CA3 pyramidal neurons is mediated by Ca^2+^- and Src family kinase-dependent endocytosis, which is facilitated by Zn^2+^-mediated suppression of tyrosine phosphatase (Eom et al., 2019; Hyun et al., 2013). The surface expression of Kv4.2 and endogenous A-type K^+^ currents in hippocampal neurons were increased by the introduction of constitutively active CaMKII (Varga et al., 2004). Taken together, these and our results suggest that the phosphorylation-dependent surface localization of Kv channels, rather than their expression, might be an important regulatory mode to modulate neuronal activity. However, we found that the levels of surface Kv4.2 proteins were not altered by the CB levels (Figs 5A, B). Further studies are required to clarify the difference between CaMKII-dependent regulation of Kv4.1 and Kv4.2. It is possible that the increase in CaMKII activity by CB expression is not sufficient to increase Kv4.2 phosphorylation to promote surface trafficking. We performed experiments without applying specific stimuli that are usually used in synaptic plasticity experiments to promote Ca^2+^ influx, while previous studies mainly focused on the role of Ca^2+^-dependent kinases and phosphatases in activity-dependent regulation. Therefore, our results represent the impact of the balance between Ca^2+^-dependent kinases and phosphatases on the phosphorylation states of Kv4.1 at resting Ca^2+^ levels. It remains to be investigated how the Kv4.1 phosphorylation state is changed upon stimulation, contributing to the activity-dependent regulation of GC excitability.

It has been established that CaMKII is dysregulated in the hippocampus of patients with AD (Ghosh and Giese, 2015). Key findings include reduced autophosphorylation of CaMKII in the hippocampus and frontal cortex of patients with AD (Amada et al., 2005), alterations in the subcellular localization of autophosphorylated CaMKII in the CA3 and DG of the AD brain (Reese et al., 2011), and inhibition of CaMKII activation by amyloid β (Zhao et al., 2004). However, it is still unclear how this dysregulation occurs. In the present study, we showed that Ca^2+^ buffering capacity is severely reduced in GCs from Tg2576 mice (Fig. 7), which we previously reported had impaired pattern separation at an early stage (Kim et al., 2021). Considering that CaMKII activation is inhibited by CB depletion (Fig. 5), reduced Ca^2+^ buffering capacity in Tg2576 GCs may contribute, at least in part, to dysregulated CaMKII signaling in AD. It is interesting to note that mice lacking CaMKII or calcineurin exhibit multiple abnormal behaviors related to schizophrenia (Miyakawa et al., 2003; Yamasaki et al., 2008).

These results together with the present study support the view that Ca^2+^ buffering deficits are causally related to hyperexcitable DG and cognitive deficits. However, abnormal excitability and CB depletion in the DG were also reported in CaMKII-deficient mice (Yamasaki et al., 2008), suggesting the possibility that CB expression is regulated by CaMKII. It is possible that CaMKII activity is regulated by Ca^2+^ and activated CaMKII, which in turn regulates various mechanisms that regulate Ca^2+^ homeostasis, including CB expression. Close interrelationships between Ca^2+^ homeostasis and the balance between excitation and inhibition may be the most important features underlying normal brain functions. Considering the presence of many Ca^2+^ buffers in neuronal cells, future studies should examine the role of different Ca^2+^ buffers in this relationship.

## Acknowledgments

This research was supported by the National Research Foundation grants from the Korean Ministry of Science and ICT (2017R1A2B2010186 and 2020R1A2B5B02002070 to W.-K. Ho)

## Author Contributions

Kim KR, Jeong HJ, Kim Y, Lee SY, and Kim HJ conducted the experiments. Kim Y, Lee SH, Cho H, Kang JS, and Ho WK designed the experiments and wrote the paper.

## Materials and Methods

### Preparation of brain slices

Brain slices were prepared from calbindin-D_28k_ knock-out (CBKO) mice (genetic background C57BL6/J), and control C57BL6/J mice aged from 1 to 2 months old. CBKO mice were kindly provided by Dr. Schwaller (Fribourg Univ. Switzerland). Average ages of CBKO and control C57BL6/J mice used in the present study were 5.6 (*n* = 36), and 5.7 week (*n* = 10), respectively. Experiments for dentate GCs were mostly conducted using mice at postnatal week (PW) 4 to PW 7, while experiments for CA1-PCs were conducted using mice at PW 3 to PW 4. Mice were killed by decapitation after being anesthetized with isoflurane, and the whole brain was immediately removed from the skull and chilled in artificial cerebrospinal fluid (aCSF) at 4 °C. Transverse hippocampal slices (350 μm thick) were prepared using a vibratome (VT1200S, Leica, Germany). Slices were incubated at 35 °C for 30 min and thereafter maintained at 32 °C until in situ slice patch recordings and fluorescence microscopy. All experiments procedures were conducted in accordance with the guide lines of University Committee on Animal Resource in Seoul National University (Approval No. SNU-090115-7).

### Electrophysiological analysis for excitability and K^+^ currents

Hippocampal GCs of DG were visualized using an upright microscope equipped with differential interference contrast (DIC) optics (BX51WI, Olympus, Japan). Electrophysiological recordings were made by the whole cell clamp technique with EPC-8 amplifier (HEKA, Lambrecht, Germany). Experiments were performed at 32 ± 1 °C. After break-in, we waited 5 min to stabilize neurons. The perfusion rate of bathing solution and the volume of the recording chamber for slices were 2.2 ml/min and 1.2 ml, respectively. Patch pipettes with a tip resistance of 3–4 MΩ were used. The series resistance (R_s_) after establishing whole-cell configuration was between 10 and 15 MΩ. The pipette solution contained (in mM): 143 K-gluconate, 7 KCl, 15 HEPES, 4 MgATP, 0.3 NaGTP, 4 Na-ascorbate, and 0.1 EGTA/or 10 BAPTA with the pH adjusted to 7.3 with KOH. For the antibody-blocking experiments, patch pipettes were dipped into an internal solution and then back-filled with the internal solution containing the antibody of Kv4.1 at a concentration of 0.3 μg/ml. The bath solution (or aCSF) for the control experiments contained the followings (in mM): 125 NaCl, 25 NaHCO_3_, 2.5 KCl, 1.25 NaH_2_PO_4_, 2 CaCl_2_, 1 MgCl_2_, 20 glucose, 1.2 pyruvate and 0.4 Na-ascorbate, pH 7.4 when saturated with carbogen (95% O_2_ and 5% CO_2_). In all bath solutions, 20 μM bicuculine and 10 μM CNQX were included to block the synaptic inputs. In voltage clamp experiments to record K^+^ currents, we added TTX (0.5 μM), CdCl_2_ (300 μM) and NiCl_2_ (500 μM) to block Na^+^ and Ca^2+^ channels, and membrane potentials were depolarized to a maximum of 30 mV for 1 s by 10 mV increments from the holding potential of −70 mV. In current clamp experiments to analyze neuronal excitability, the following parameters were measured: (1) the resting membrane potential (RMP), (2) the input resistance (R_in_, membrane potential changes (V) for given hyperpolarizing current (−35 pA, 600 ms) input), (3) F-I curve (firing frequencies (F) against the amplitude of injected currents (I), for DG; 100 pA to 600 pA, 100 pA increment, 1 s duration, for CA1 pyramidal cells (CA1 PCs); 50 pA to 250 pA, 50 pA increment, 1 s duration), (4) AP onset time at 300 pA (the delay from the start of 300 pA injection to the beginning of the upstroke phase of the 1^st^ evoked AP). All chemicals were obtained from Sigma (St. Louis, MO, USA), except CNQX, bicuculline, and TTX from abcam Biochemicals (Cambridge, UK).

### Cytosolic Ca^2+^ measurement and estimation of calcium binding ratios

Cytosolic [Ca^2+^] was measured from fluorescence images of a hippocampal GCs in the slice loaded with fura-2 (pentapotassium salt), using a polychromatic light source (xenon-lamp based, Polychrome,Martinsried, Germany), which was coupled into the epi-illumination port of an upright microscope (BX51, Olympus, Japan) via a quartz light guide and an UV condenser, and an air-cooled slow-scan CCD camera (SensiCam, PCO, Kelheim, Germany). The monochromator and the CCD camera were controlled by a PC and ITC18, running a custom-made software programmed with Microsoft Visual C^++^ (version 6.0). Details for calibration method were described previously (Lee et al., 2012).

To estimate the endogenous Ca^2+^ binding ratio (κ_E_), we applied short depolarizing pulses (from −70 mV to 0 mV, 40 ms duration) every 20–60 s in order to evoke [Ca^2+^] transients in the mature dentate GCs. Normally, κ_E_ were estimated according to the single compartment and linear approximation model (Neher, 1998; Neher and Augustine, 1992). When two kinds of Ca^2+^ buffer, Ca^2+^-indicator dye (B) and endogenous Ca^2+^ buffer (E), exist in the compartment, increments of total and free calcium have the following relationship:

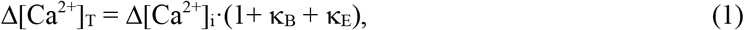

where κ_B_ and κ_E_ are calcium binding ratios of B and S, respectively. Ca^2+^ transients (Δ[Ca^2+^](t), CaTs) following short pulses of Ca^2+^ influx can be described by the following equation:

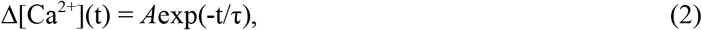

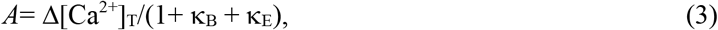

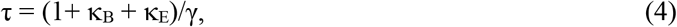

where *A* is the initial amplitude, d[Ca^2+^]_T_ is the total intracellular Ca^2+^ increase evoked by the influx, γ is the Ca^2+^ extrusion rate and κ_B_ is the Ca^2+^ binding ration of the Ca^2+^ indicator dye (fura-2). Calcium binding ratio of a buffer, X, is defined by

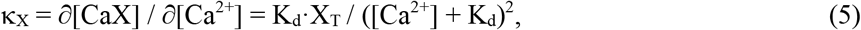

where X_T_ and K_d_ are total concentration of X and the dissociation constant of X for Ca^2+^, respectively (Neher and Augustine, 1992). When κ_X_ is not constant over the dynamic range of [Ca^2+^] in a Ca^2+^ transient, incremental Ca^2+^-binding ratio, κ′_X_, should be used, which is defined as

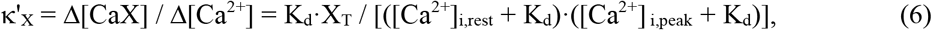

where [Ca^2+^]_i,rest_ and [Ca^2+^]_i,peak_ represent [Ca^2+^]_i_ value before and at the peak perturbation, [X]_T_ is the total concentration of Ca^2+^ buffer X, and K_d_ is the Ca^2+^ dissociation constant of X (Neher and Augustine, 1992). According to equation (3) and (4), plotting *A* and τ values measured at different level of κ_B_ provides two estimations of κ_E_: one is obtained from the *x*-intercept of the straight line fit to the plot of τ vs. κ_B_, the other from a line fit to the plot of *A^−1^* vs. κ_B_. These two estimates will be referred to as κ_E_.

### Cell culture and constructs

HT22 (Sigma-Aldrich #SCC129) and HEK293T (ATCC CRL-3216) cells were cultured as previously described(Cesarini et al., 2018; Pyun et al., 2018). HT22 and HEK293T cells were culture in DMEM medium containing 10% fetal bovine serum (FBS, Gibco) and transiently transfected with a combination of DNA plasmids by using lipofectamine 3000 (Invitrogen). To examine CB knockdown effects, HT22 cells were infected with lentivirus carrying the control scrambled-shRNA (5’-CCGGCAACAAGATGAAGAGCACCAACTCGAGTTGGTGCTCTTCATCTTGTTGTTTTTG-3’) or CB-specific shRNA (5’-CCGGGATTGGAGCTATCACCGGAAACTCGAGTTTCCGGTGATAGCTCCAATCTTTTTG-3’) with polybrene (Sigma) for two days. To induce differentiation, cells at approximately 70-80% confluence were switched to neurobasal medium (Gibco) containing N2, B27, glutamax, 5ng/ml BDNF and 50ng/ml NGF (Thermo fisher). Constructs used in this study are as following: pCI-neo, pCI-neo-CB, Kv4.1-GFP (Origene, MG220056) and GFP-C1-CaMKIIα (addgene, 21226). To examine the role of Kv4.1 phosphorylation, the serine-alanine mutation in Kv4.1 at serine residue 265, 555 or 568 was introduced by using a mutagenesis kit (Stratagene).

The Kv4.1 currents from HEK293T cells were measured with the whole cell patch-clamp technique. Voltage clamp was performed by using an EPC-10 amplifier (HEKA Instrument, Germany) and filtered at 10 kHz. The patch pipettes (World Precision Instruments, Inc., USA) were made by a Narishige puller (PP-830, Narishige Co, Ltd., Japan). The patch pipettes used had a resistance of 2–3 MΩ when filled with the pipette solutions listed in the next sections. All recordings were carried out at room temperature. The normal external solution for HEK293T cell recording was as follows (in mM): 143 NaCl, 5.4 KCl, 5 HEPES, 0.5 NaH2PO4, 11.1 glucose, 0.5 MgCl2, 1.8 CaCl2, pH 7.4 adjusted with NaOH. The pipette solution was as follows (in mM): 135 K-aspartate, 2 MgCl2, 3 EGTA, 1 CaCl2, 4 Mg-ATP, 0.1 Na-GTP, 10 HEPES, pH 7.4 adjusted with KOH. Currents were analyzed and fitted using Patch master (HEKA Instrument) and Origin 6.1 (Originlab Corp., USA) software. All values are given as mean ± SEM. Current-voltage (I/V) relations were obtained by plotting the outward current at 1-s of test pulse as a function of the test potential. Current densities (pA/pF) were obtained after normalization to cell surface area calculated by Patch master.

### Protein analysis and Surface biotinylation

Western blot analysis was performed as previously described (Jeong et al., 2019; Kim et al., 2020). Briefly, cells were lysed in RIPA lysis buffer (iNtRON) containing complete protease inhibitor cocktail (Roche), followed by SDS-PAGE and incubation with primary and secondary antibodies. Primary antibodies used are Calbindin, pCaMKIIα, CaMKIIα (Cell signaling), Kv4.1, Kv4.2 (Alomone lab), GFP (Abcam) and β-tubulin (Sigma).

Immunoprecipitation were performed as previously described (Choi et al., 2019). Briefly, precleared cell extracts were incubated with phospho-Serine antibodies (Santacruz) overnight at 4 °C, followed by incubation with protein G-agarose beads (Roche) for 1 hour. Subsequently beads were washed three times with cell extraction buffer and precipitates were subjected to western blotting.

Surface biotinylation analysis was carried out as previously described (Bae et al., 2020). In brief, the surface proteins of HT22 cells or mouse dentate gyrus were biotinylated by exposure to 1mg/ml NHS-LC-biotin (Thermo) for 30-60 minutes at 4°C. After quenching with 100mM glycine, cells or dentate gyrus were lysed and sonicated in RIPA lysis buffer (iNtRON) with proteinase inhibitor (Roche), followed by incubation with streptavidine-agarose beads (Pierce) and western botting. Immunoreactivity was determined by enhanced chemiluminescence (GE Healthcare). The images were captured using a bioimaging analyzer (LAS-3000; Fuji, Tokyo, Japan) and analyzed with a Multi-Gauge program (Fuji, Tokyo, Japan).

### Behavior Analysis

Contextual fear conditioning was performed with male mice between 15 and 16 weeks of age in 8 CBKO mice. We modified the protocol of McHugh et al (2007) to conduct the contextual fear discrimination task. Mice were trained to discriminate between two similar contexts, A and B, through repeated experience in each context. Context A (conditioning context) is a chamber (18 cm wide × 18 cm long × 30 cm high; H10-11M-TC; Coulbourn Instruments 5583, PA 18052, USA) consisting of a metal grid floor, aluminium side walls, and a clear Plexiglass front door and back wall. Context A was indirectly illuminated with a 12 W light bulb. The features of Context B (safe context) were the same as Context A, except for a unique scent (1% acetic acid), dimmer light (50% of A), and a sloped floor by 15° angle. Each context was cleaned with 70% ethanol before the animals were placed. On the first 3 days (contextual fear acquisition), the mice were placed in Context A for 3 min for exploring the environment, and then received a single foot shock (0.75 mA, for 2 s). The mice were returned to their home cage 1 min after the shock. On day 4 – 5, mice of each genotype were divided into two groups; one group visited Context A on Day 4 and Context B on Day 5, while the other group visited the Context B on Day 4 and Context A on Day 5. On day 4 – 5 (generalization), neither group received a shock in Context A and B, and freezing level was measured for 5 min only in Context A. We defined freezing behavior as behavioral immobility except for respiration movement (McNaughton & Nadel, 1990). We observed video image for 2 s bouts every 10 s (18 or 30 observation bouts for 3 min or 5 min recording time) and counted the number of 2 s bouts during which the mouse displayed freezing behavior (referred to as a freezing score). The percentage of freezing was calculated by dividing the freezing score with the total number of observation bouts (18 or 30). The mice were subsequently trained to discriminate these two contexts by visiting the two contexts daily for 8 days (from day 6 to 13, discrimination task). The mice always received a footshock 3 min after being placed in Context A but not B (Fig. 7C). Discrimination ratios were calculated according to F_A_ / (F_A_ + F_B_), where F_A_ and F_B_ are freezing scores in Contexts A and B, respectively. All experiments and analyses were performed blind to the mice genotype.

For one trial contextual fear conditioning, experiment was performed with male mice between 14 and 19 weeks of age (8 control mice and 8 CBKO mice) in a pair of very distinct contexts (A and C). The aforementioned Context A was used as the conditioning context. The distinct context (Context C) is a white acrylic blind end cylinder (15 cm in diameter, 18 cm in height, and 0.5 cm in thickness) standing vertically on the metal grid floor of Context A, and the bottom inside the cylinder was covered with cage bedding, on which mice were placed. The chamber and cylinder were cleaned using 70% ethanol between runs. On day 1 (acclimation), mice were placed in Context A and then placed them in Context C an hour later. Mice were allowed to freely explore in both contexts for 5 min. On day 2 (conditioning), all group of mice were place in Context A and receive a single foot shock (0.75 mA, for 2 s) 3 min later. Mice were left in the Context A for 1 min after the shock. On day 3 (assessment), mice were separated into two groups; mice of each group were placed in Context A or C for 3 min without a foot shock, during which the freezing score was measured.

Open field exploration test was used to assess locomotor activity (Kim et al., 2012). Open field exploration was performed with male mice 16 weeks of age (6 control mice and 6 CBKO mice). Mice were handled before open field exploration behavior test (OFT). Mice were picked up by the tail and held and without restraint in the palm of the hand for 15 min/day for 4 days. Following a handling, they were placed in a holding cage until all mice were handled. They were then returned to their home cage. During the mice handling, handlers wore latex gloves and changed the glove between the mice. After handling day, we were executed OFT test. we modified the protocol of Kim et al (2012) to conduct the OFT test. The Open field box was made of white plastic (40 × 40 × 40 cm) under diffuse lighting and white noise and the open field was divided into a center zone (center, 20 × 20 cm and an outer field. Individual mice were placed in the center zone and the path the animal were recorded with a video camera. During a 20 min observation period, the total distance traveled for 20 min and time spent in the center zone for initial 5 min were analyzed using the program EthoVision XT (Noldus, Virginia, USA)

### Statistical analysis

Data were analyzed with IgorPro (Version 6; Wave-Metrics, Lake Oswego, USA) and are presented as the mean ± S.E.M. with the number of cells or mice (n) used in each experiment. The statistical significances were evaluated using Student’s t-test, and the level of significance was indicated by the number of marks (*p< 0.05; **p < 0.01; ***p < 0.001). p *>* 0.05 was regarded as not significantly different (N.S.). Data from behavioral studies were analyzed using Igor Pro and Origin (Version 8; Microcal, Northampton, USA). Comparison between multifactorial statistical data was made using two-way analysis of variance (ANOVA). The differences in time-dependent changes of behavioral parameters between the two genotypes were evaluated using two-way repeated measures ANOVA.

---

**Fig. S1.**
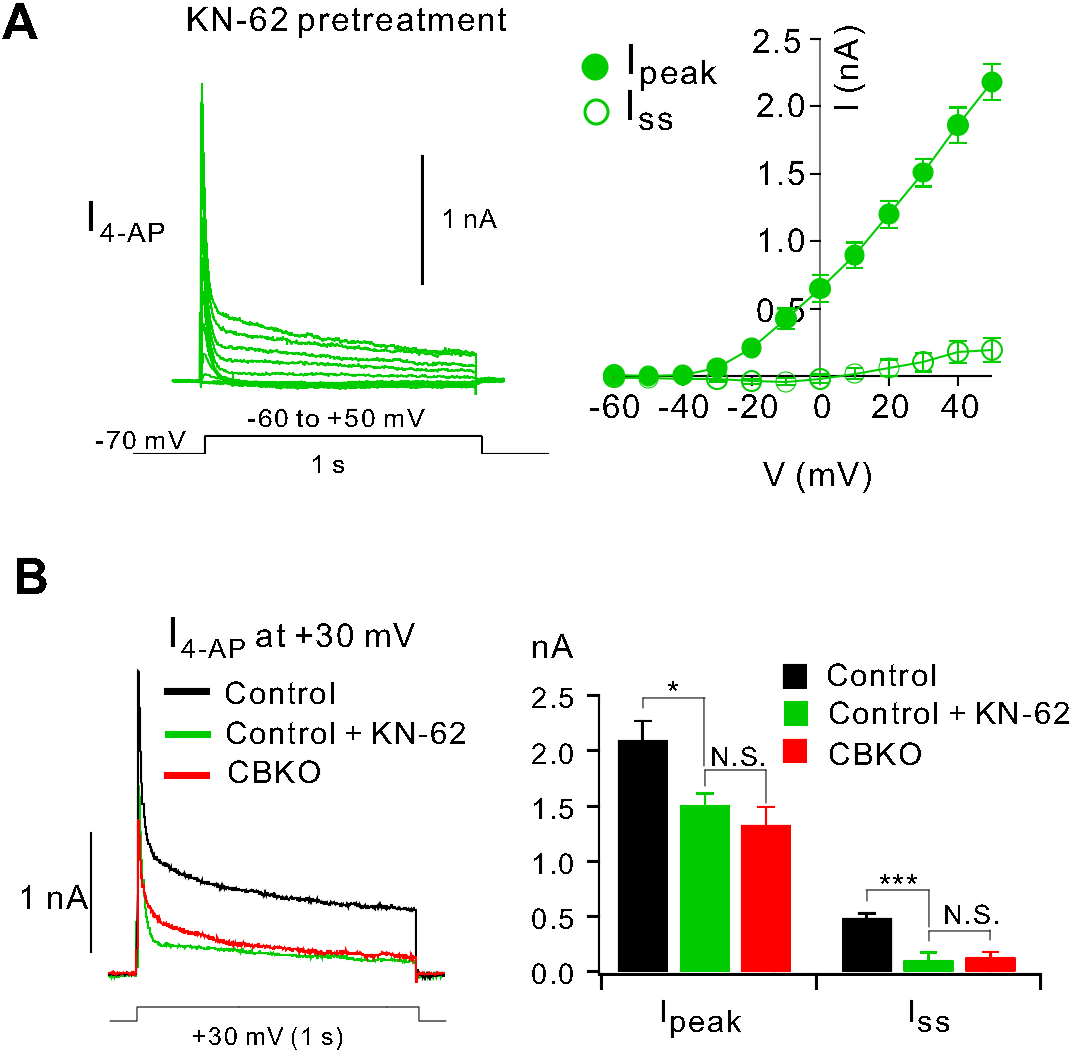
Reduced 4-AP sensitive K^+^ currents by KN-93. ***A,*** Average I_4-AP_ traces obtained from control GCs pretreated with KN-62 (*n* = *6*) for more than 20 min (left) and plots for I_peak_ and I_ss_ against the voltage (right). ***B,** Left,* I_4-AP_ trace at +30 mV obtained in the presence of KN-62 in control GCs (green) is compared with that in CBKO (red) and in control (black) shown in Fig. 2Cb. *Right,* Bar graphs for I_peak_ and I_ss_ in each condition. Data are represented as mean ± SEM. *p < 0.05, ***p < 0.001, N.S. not significant by Student’s t-test

**Fig. S2.**
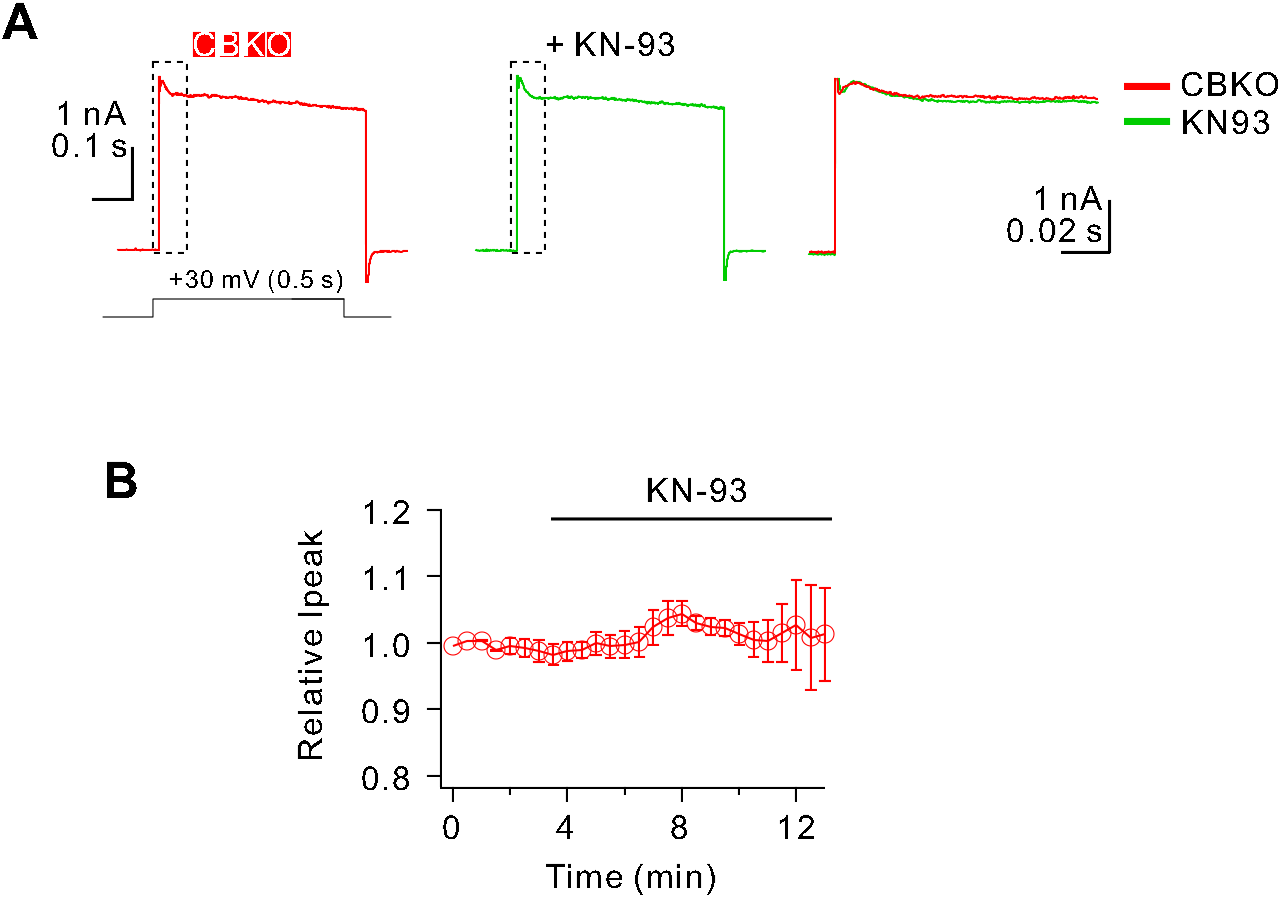
K^+^ current in CBKO GCs is not affected by N-93. ***A,*** Whole-cell K^+^ currents at + 30 mV in control GCs before (black) and after KN-93 application (green). ***B,*** Time course of the changes in K^+^ current amplitude after KN93 application (average from 7 cells).

